# Host-microbiome mutualisms emerge from community interactions among microbes

**DOI:** 10.1101/2023.11.14.567078

**Authors:** Jason R. Laurich, Emma Lash, Megan E. Frederickson

**Affiliations:** Department of Ecology & Evolutionary Biology, University of Toronto, Toronto, ON, Canada; Department of Molecular Genetics, University of Toronto, Toronto, ON, Canada

## Abstract

Microbiomes often benefit plants, conferring resistance to pathogens, improving stress tolerance, or promoting plant growth. As potential plant mutualists, however, microbiomes are not a single organism but a community of species with complex interactions among microbial taxa and between microbes and their shared host. The nature of ecological interactions among microbes in the microbiome can have important consequences for the net effects of microbiomes on hosts. Here, we compared the effects of individual microbial strains and 10-strain synthetic communities on microbial productivity and host growth using the common duckweed *Lemna minor* and a synthetic, simplified version of its native microbiome. Except for *Pseudomonas protegens*, which was a mutualist when tested alone, all of the single strains we tested were commensals on hosts, benefiting from plant presence but not increasing host growth relative to uninoculated controls. However, 10-strain synthetic microbial communities increased both microbial productivity and duckweed growth more than the average single-strain inoculation and uninoculated controls, meaning that host-microbiome mutualisms can emerge from community interactions among microbes on hosts. The effects of community inoculation were sub-additive, suggesting at least some competition among microbes in the duckweed microbiome. We also investigated the relationship between *L. minor* fitness and that of its microbes, providing some of the first empirical estimates of broad fitness alignment between plants and members of their microbiomes; hosts grew faster with more productive microbes or microbiomes.

## Introduction

Plants and animals harbour diverse microbiota that often affect the phenotypes or fitness of their hosts. The microbes in these microbiomes interact with one another, just like plants and animals do in more familiar ecosystems such as grasslands or rainforests. Microbes living together in or on hosts can compete for resources, exploit other microbes in interactions akin to predation or parasitism, or cooperate in mutualisms. However, the relative importance of these interaction types (i.e., competition, exploitation, or mutualism) in microbial communities is the subject of debate (Palmer and Foster 2022), and microbiome science is only beginning to interrogate the consequences of microbial interactions within the microbiome for their combined effects on hosts (Hassani et al. 2018).

Several studies have assessed the frequency of competition, exploitation, and mutualism among microbes in communities (Foster and Bell 2012; Gould et al. 2018; Kehe et al. 2021; Ortiz et al. 2021; Palmer and Foster 2022). One tested many possible combinations of 72 bacteria isolated from rainwater pools in tree holes and found that most bacteria grow better alone than with other microbes (Foster and Bell 2012), suggesting that competition prevails among culturable bacteria. However, when Kehe et al. (2021) used an ultrahigh-throughput platform to test over 180,000 combinations of 20 soil bacteria, they found more positive interactions than expected; about 40% of bacteria grew better with other bacteria than alone, although most of these interactions were exploitative, not mutualistic. Recently, Palmer and Foster (2022) synthesized the available evidence from multiple studies of microbial communities and concluded that “negative interactions prevail, and cooperation, where both species benefit, is typically rare.”

Competition among microbes in the microbiome may benefit hosts, if host pathogens are competitively suppressed or excluded by other microbes (Wang et al., 2022). Pathogen suppression is a primary benefit of plant microbiomes that is often mediated by a small subset of strains (e.g., Hu et al. 2016; Ishizawa et al. 2019; Barelli et al. 2020; Löser et al. 2021). If pathogen suppression emerges as a consequence of competition among strains, then internal antagonism in microbiomes has the potential to result in overall microbiome cooperation with hosts. Such dynamics could also explain why more diverse microbiomes can provide greater pathogen-suppression benefits to their hosts (Hu et al. 2016). Consistent with classical ecological theory, more diverse microbiomes may be more likely to contain strains that suppress pathogens through competitive dominance (e.g., the sampling effect, Tilman et al. 1997a). This raises the possibility that mutualistic outcomes between certain pathogen-suppressing microbes and hosts may be more visible with increasing microbiome complexity, precisely because more diverse communities are more likely to contain the very pathogens whose inhibition demonstrates their positive effects. More generally, diversity within the microbiome can increase the overall productivity and ecosystem services provided by plant microbes if diverse microbiomes are more likely to contain ‘keystone’ microbes that have disproportionately large direct effects on host phenotypes, or strongly shape microbial community composition (e.g., Coyte et al. 2015; Tang et al. 2015; Agler et al. 2016; Hu et al. 2016; Zhang et al. 2017; Melnyk et al. 2019; Löser et al. 2021; Tan et al. 2021).

However, not all interactions among microbes are competitive, and we might expect more positive interactions among microbes in the host environment than in free-living communities. Few of the studies highlighted by Palmer and Foster (2022) grew microbes in association with a host (but see Gould et al. 2018; Ortiz et al. 2021), and even fewer simultaneously compared single and multiple strains of microbes in terms of their effects on both microbial and host growth. The presence of a host has the potential to change the nature of microbe-microbe interactions within a community, potentially in favor of more positive interaction outcomes. Even if two microbes compete for resources, if one microbe promotes host growth in a way that generates increased supply of host rewards to the microbiome as a whole, then other microbes will benefit from its presence. Plant-derived organic carbon is likely one such shared reward for microbes; plants secrete up to 44% of their fixed carbon as root exudates (Bais et al. 2006), resulting in microbial densities in the rhizosphere far in excess of microbial densities in surrounding environments (e.g., Coler and Gunner 1969).

Several studies have found that more diverse bacterial communities are more productive or provide greater ecosystem services (e.g., Bell et al. 2005; Delgado-Baquerizo et al. 2016; Hu et al. 2016; Maron et al. 2018; O’Brien et al. 2022), a microbial version of the classic biodiversity-ecosystem function (BEF) relationship often documented for plant communities (Tilman et al. 1997b). Positive BEF relationships can arise through competitive dynamics and sampling or portfolio effects, but facilitation among microbes likely also contributes (Hassani et al. 2018). Many bacterial metabolites are leaky, diffusing across cell membranes where they can be taken up and consumed by other community members. This dynamic can lead to the evolution of cross-feeding or nutritional dependence among bacteria, and thereby increasing productivity in more diverse microbial communities (Hoek et al. 2016; D’Souza et al. 2018). In addition, indirect ecosystem services, such as the degradation of antibiotics or toxins, or biofilm formation, can be strongly selected for in certain community members through market-like dynamics that redound to the benefit of all strains (Piccardi et al. 2019; Adkins-Jablonsky et al. 2021). The exchange of energetically expensive bacterial metabolites such as amino acids or carbohydrates, and the partitioning of ecosystem service provisioning can lead to ‘Black Queen’ dynamics, wherein selection favours the down-regulation or outright loss of expensive metabolic pathways provided by other community members (Morris et al. 2012; Morris 2015). Selection for this metabolic stream-lining can be very strong (D’Souza et al. 2014), and can result in the evolution of ecological specialization and nutritional inter-dependence that increases the overall productivity of communities (D’Souza et al. 2014; Hoek et al. 2016; D’Souza et al. 2018; Pacheco et al. 2019). In host-microbiome interactions, the production of functionally distinct rewards by different community members and the evolution of niche complementarity among them is also expected to increase the sum of benefits obtained by their hosts (Afkhami et al. 2014, 2021).

Whether microbes benefit one another indirectly by promoting the growth of their shared host depends on what kind of benefits microbes confer to hosts and whether those benefits feed back to all microbes living on a host (i.e., as a ‘public good’) or to only one or a few strains. In plants, in addition to suppressing pathogens (e.g., Hu et al. 2016; Ishizawa et al. 2019; Barelli et al. 2020; Kalachova et al. 2022), microbes can confer resilience against environmental stressors such as elevated salinity (Yuan et al. 2016; Mueller et al. 2021), drought (Marasco et al. 2012), or flooding (Bal and Adhya 2021), or even degrade or detoxify detrimental pollutants such as chromium, arsenic, or phenols (Tang et al. 2015; Zhang et al. 2017; Radulović et al. 2020). Plant microbiomes can also promote plant growth by fixing nitrogen (Duong and Tiedje 1985; Fonseca et al. 2018), solubilizing phosphates (Ishizawa et al. 2017), or producing plant-growth promoting hormones or compounds such as indoles and auxins (Gilbert et al. 2018; Tan et al. 2021). However, whether these benefits of microbes to hosts feed back to benefit the microbes themselves remains an open empirical question in most systems, because few studies measure the benefits of host association to microbes or the extent of fitness alignment or conflict between host and microbial partners (Douglas and Werren 2016; Mushegian and Ebert 2016). Fitness feedbacks between hosts and symbionts determine how cooperation evolves between species (Trivers 1971), and the evolution of genuinely mutualistic interactions between plants and microbes is no less dependent on such factors than other relationships (Hawkes et al. 2020; O’Brien et al. 2021).

The only plant-microbe interaction in which fitness alignment or conflict has received substantial attention is the legume-rhizobium mutualism, in which legumes host rhizobacteria in root nodules where they exchange fixed carbon for fixed nitrogen. Inoculations of rhizobia strains onto legumes have revealed mostly positive fitness correlations between partners (Friesen 2012; Frederickson 2017), implying that natural selection generally favors the evolution of more beneficial rhizobia. Indeed, recent evolution experiments with rhizobia have directly observed the evolution of greater host benefits in real time (Batstone et al. 2020). In contrast, whether plant-microbe fitness correlations are positive, negative, or neutral in other systems is a largely open question (but see O’Brien et al. (2020a)), meaning we have a limited understanding of whether selection favors more or less beneficial microbes in symbioses beyond legumes and rhizobia. That plants often benefit substantially from their microbiomes is well documented (e.g., Marasco et al. 2012; Mueller et al. 2021; Kalachova et al. 2022), and many plants invest heavily in the sort of reciprocal exchange of nutrients that fuel mutualistic interactions through rewards such as root exudates (Bais et al. 2006). However, we would also expect to find that some microbes in plant microbiomes are pathogenic and proliferate rapidly by over-exploiting plants and reducing plant fitness (Melnyk et al. 2019). Furthermore, fitness correlations measured in legumes and rhizobia generally involve comparing the performance of many closely related rhizobia strains on hosts (i.e., phenotyping many isolates of the same rhizobium species), while plant microbiomes are highly diverse with many microbial lineages competing for host rewards. Whether natural selection favors the most beneficial microbes in diverse plant microbiomes, or whether microbial fitness is largely uncoupled from plant benefits, deserves greater empirical attention in plant-microbiome interactions.

Here, we leveraged the relationship between the common duckweed *Lemna minor* and its microbiome to investigate several fundamental questions pertaining to the ecology and evolution of plant-microbiome interactions. Duckweeds (Lemnaceae) are the world’s fastest growing and smallest angiosperms (Bog et al. 2019). Their rapid growth rates, coupled with their nearly entirely clonal reproduction through the budding of fronds (Landolt 1986; Ho 2018), facilitates measurements of host fitness at high replication in a laboratory setting (Kose et al. 2022). In this study, we compared the effects of single microbial strains and 10-strain synthetic microbial communities inoculated onto sterilized *L. minor* plants. We measured the effects of these treatments on duckweed fitness and the productivity of *L. minor* microbiomes, and quantified the degree of fitness alignment between *L. minor* and its microbes. Specifically, we sought to address the following questions: (1) How do interactions among microbes affect microbial productivity in the host versus free-living environment? (2) How do microbiome diversity and microbe-microbe interactions affect the benefits microbiomes provide to their hosts? And (3), how aligned are the fitness interests of *L. minor* and its microbes?

## Materials and methods

### Collection and culturing of *Lemna minor* and its microbiome

*Lemna minor* is a small, floating, aquatic macrophyte with a worldwide distribution in temperate zones. Like other duckweeds, *L. minor* exhibits a highly reduced morphology and simple life history (Bog et al. 2019). Plants consist of 1-4 fronds ranging in length from 1 to 8 mm, with a single adventitious root per frond (Landolt 1986). While *L. minor* is capable of sexual reproduction, flowering is extremely rare in *L. minor* (Pieterse 2013), and populations have little segregating genetic diversity due to high rates of clonal reproduction through frond budding (Landolt 1986; Cole and Voskuil 1996; Ho 2018). *Lemna minor* boasts an extremely high growth rate, doubling approximately every four days (Hossell and Baker. 1979), and can establish dense, dominant vegetative mats in ponds and other slow-moving bodies of water (Keddy 1976; Driever et al. 2005). Distinguishing *L. minor* from other closely related duckweeds can be difficult as a result of their highly simplified morphology, and genetic markers suggest that plants identified as *L. minor* may sometimes be *L*. × *japonica*, a hybrid of *L. minor* and *L. turionifera* (Braglia et al. 2021; Volkova et al. 2023). Nonetheless, in keeping with previous studies (O’Brien et al. 2020a,b), we refer to the duckweeds used in these experiments as *L. minor*.

We collected *L. minor* plants from two locations in the Greater Toronto area in the summer of 2017: Churchill Marsh (43.77°N, 80.02°W) and Wellspring Pond (43.48°N, 79.72°W). Immediately after collecting live plants in the field, we cultured components of the *Lemna minor* microbiome by crushing the tissue of approximately six fronds per population and spreading this mixture onto yeast mannitol agar plates. We cultured plates at 29 °C for 5 days to generate a master plate containing a diversity of culturable strains present in the duckweed microbiome. These plates capture only a subset of the bacterial strains present in the duckweed microbiome, displaying an order of magnitude less diversity than plants in the field (O’Brien et al., 2020a,b). The microbes we cultured are biased towards epiphytic bacterial strains, and under-represent the diverse bacterial endophytes, fungi, and diatoms present in *L. minor* (Rejmankova et al. 1986; Goldsborough 2004; Gilbert et al. 2018; Acosta et al. 2020; O’Brien et al. 2020a,b; Tan et al. 2021). Nevertheless, they represent many of the most abundant taxa present in the *L. minor* microbiome (O’Brien et al. 2020a,b), and many of these bacteria affect duckweed growth and phenotypes (Duong and Tiedje 1985; Tang et al. 2015; Ishizawa et al. 2017; O’Brien et al. 2020a,b; Tan et al. 2021). We isolated single bacterial strains from these master plates by serially re-streaking and plating colonies until the colonies had a single phenotype. We then grew liquid cultures of individual bacteria by inoculating liquid yeast mannitol media with cells taken from a single colony. We placed these liquid cultures in a shaking incubator at 29 °C for 5 days to generate experimental inocula. We then used a NovoCyte Flow Cytometer (ACEA Biosciences, Inc., San Diego, CA, USA) to measure the concentration of bacterial cultures in liquid media before diluting to 10^5^ cells per 50 µL of inoculum.

To generate isoclonal lines of *L. minor*, we transferred duckweed fronds from each population to Krazčič growth media (Krazčič et al. 1995) in 500 mL Mason jars. We maintained stock cultures in an environmental chamber set to cycle between a 16 hour period at 23°C and 150 µmol/m^2^ light followed by 8 hours of darkness at 18°C. We then placed individual fronds in sterile media; all plants used in subsequent analyses and experiments are the clonal descendants of a single frond per population. Once our isogenic cultures reached high density, we sterilized duckweed fronds by vortexing plants twice in phosphate-buffered saline for 5 minutes before immersing them in 1% sodium hypochlorite bleach for one minute. We then thoroughly rinsed bleached fronds in autoclaved distilled water and transferred plants to sterile growth media. This sterilization process disrupts and simplifies the natural microbiome of *L. minor*, and is particularly effective at sterilizing epiphytic bacteria associated with duckweed fronds and roots, but does not always entirely remove endophytic species (Gilbert et al. 2018; Tan et al. 2021). We tested the effective sterility of *L. minor* fronds roughly one week later by placing them in growth media enriched with 5 g/L sucrose and 1g/L yeast extract.

### Sequencing of bacterial strains

We extracted bacterial DNA from liquid cultures using a Polyzyme digestion protocol (Laurich et al., 2023). We digested cells in MetaPolyzyme Multilytic Enzyme mix (Sigma) and lysed cells in Puregene Cell Lysis solution (Qiagen). After extracting DNA, we performed PCR reactions using universal bacterial 16S rDNA primers (Weisburg et al. 1991). We then purified our PCR products with a QIAquick PCR Purification Kit (Qiagen), and sent purified DNA to the Centre for the Analysis of Genome Evolution and Function at the University of Toronto for Sanger sequencing on an Applied Biosystems 3730 DNA Analyzer using standard protocols. We identified our strains using the megablast algorithm against the NCBI 16S ribosomal RNA database for Bacteria and Archaea (Zhang et al. 2000).

### Experimental design and procedures

We inoculated sterilized *Lemna minor* plants from both field sites with individual bacterial strains or 10-strain synthetic bacterial communities. We measured how individual bacterial strains and synthetic communities (20 replicates per treatment) affected both microbial and host growth. For each population, plants were inoculated with microbes cultured from field-collected plants from that same population; i.e., we inoculated Churchill duckweeds with Churchill microbes, and Wellspring duckweeds with Wellspring microbes. Thus, the plant-microbe associations may be locally or co-adapted (O’Brien et al., 2023).

We filled 24-well plates with autoclaved, sterile culture media. After rinsing to remove all traces of enriched media from sterilized plants, we randomly assigned plants (1-3 fronds) to wells, leaving some wells empty for the no-host treatment. We then sealed plates with a gas-permeable membrane (Breathe Easier, Sigma-Aldrich). We inoculated each well with 50 µL of a randomly-assigned bacterial or control inoculum by pipetting through the gas-permeable membrane, and re-sealed plates with a second membrane upon completion (Breathe Easy, Sigma-Aldrich). The control inoculum consisted of autoclaved culture media, while inocula for single bacterial strain treatments contained approx. 10^5^ cells, and the synthetic microbiome inoculum contained approx. 10^4^ cells from each of 10 single bacterial cultures (resulting in a 10-strain community with the same final cell concentration as single-strain inocula). We included a no-host treatment in which we inoculated wells with bacteria, but did not add plants, to compare bacterial growth in the presence and absence of hosts.

After loading wells with plants, bacteria, or both, we took images of the plates, and placed them in an environmental chamber (16:8h light/dark, 23/18 °C cycle) for 10 days. At the end of the experiment, we photographed plates a second time. For all images, we used ImageJ (Schneider et al. 2012) to calculate the total surface area occupied by *Lemna minor*. After the experiment, we ran 10 µL of well fluid through our NovoCyte Flow Cytometer to determine the final concentration of bacterial cells in wells. We measured cell density (absolute cell count) and reduced noise by filtering out particulates below a certain size. We thresholded our samples based on forward and side scatter (threshold of 5000) to exclude fragments of dead cells and plant matter. While we did not use fluorescent stains to distinguish live from dead bacterial cells, we thresholded our samples conservatively to reduce noise, and our estimates, which will include live and dead bacterial cells (Gasol and Del Giorgio. 2000; Ou et al. 2017), are likely a good proxy of bacterial fitness or community size.

## Data analysis

We fit statistical models in R version 4.3.1 (R Core Team 2023). We modelled microbial cell density and duckweed growth in two ways. First, we tested whether plant or microbial productivity with a 10-strain community exceeded plant or microbial productivity in the average single-strain treatment, akin to a positive biodiversity-ecosystem function relationship. We fit linear mixed models using the ‘lme4’ and ‘lmerTest’ packages (Bates et al. 2015, Kuznetsova et al. 2017), including well location (edge or interior), plate number, and the identity of the bacterial strain nested within population as random effects. The plant growth model fit final duckweed area (in pixels) as a function of the fixed effects of initial duckweed area (in pixels) and treatment (three levels: control, single-strain inoculation, or 10-strain inoculation). For the plant growth model, we then used the ‘emmeans’ package (Lenth 2019) for pairwise comparisons among treatment means using the Tukey method. For the microbial growth model, we subset the data to only wells that received microbes, and fit final microbial cell density (log-transformed to improve the normality of residuals) as a function of the fixed effects of treatment (two levels: single-strain inoculation or 10-strain inoculation), plant presence, and the interaction between treatment and plant presence.

We also wanted to compare the effect of inoculating with a 10-strain synthetic community to the additive expectation from the single strain effects to ask whether microbes have sub-additive or synergistic effects on plant and microbial productivity (Afkhami et al., 2014). For example, when the productivity of the 10-strain community is less than the sum of the cell densities of the 10 strains grown alone, the results suggest more microbe-microbe competition than mutualism (Foster and Bell 2012). For the microbes, we modelled the cell density of each population separately, with bacterial treatment (11 levels: the 10 single strains and the 10-strain community), plant presence, and the interaction between bacterial treatment and plant presence as fixed effects and well location (edge/interior) and plate number as random effects. For plant growth, we modelled the final duckweed area (in pixels) of each population separately, with initial duckweed area (in pixels) and all 12 bacterial treatments (the 10 single strains, the 10-strain community, and control, uninoculated plants) as fixed effects, again including well location (edge/interior) and plate number as random effects. We considered the 10-strain synthetic community effect to be significantly different from the additive expectation from single strain inoculations if the 95% confidence interval of the estimated marginal mean for the 10-strain community did not overlap the sum of the model coefficients for the 10 single strains (plus the control, in the case of plant growth only). We calculated estimated marginal means and additive expectations on the raw scale to facilitate comparisons and interpretation.

Finally, to examine fitness correlations between hosts and microbes, we again used the ‘emmeans’ package (Lenth 2019) to extract the bacterial treatment means from linear mixed effects models for duckweed growth and microbial cell density. These models were fit on data standardized to a mean of 0 and standard deviation of 1 within each population, and again included the random effects of edge and plate number. Our growth models also included standardized initial pixel count as a fixed effect. As the plants used in our experiments are all clones and each of our bacterial treatments consisted of inocula developed from a single colony, variation within bacterial treatments is environmental, while variation in both microbe and host fitness among bacterial treatments is due to genetic differences among bacteria. We fit a simple linear regression between bacterial treatment means for duckweed and microbial growth to determine whether the fitness interests of microbes and duckweeds are aligned in each population.

## Results

### Bacterial strain diversity

We isolated a diversity of bacteria strains, with only *Pseudomonas protegens* shared between Churchill and Wellspring isolates. Bacteria isolated from Churchill duckweeds also included other Proteobacteria in the families Aeromonadaceae (*Aeromonas*), Acetobacteraceae (*Falsiroseomonas*), and Boseaceae (*Bosea*), as well as Bacteroidetes in the families Chitinophagaceae (*Parasediminibacterium*), Flavobacteriaceae (*Flavobacterium*), Fulvivirgaceae (*Ohtaekwangia*), and Spirosomaceae (*Arcicella*) and finally Actinobacteria in the family Microbacteriaceae (*Microbacterium*). Isolates from Wellspring duckweeds were almost all Proteobacteria, including strains in the Sphingomonadaceae (*Sphingomonas* and *Rhizorhabdus*), Rhizobiaceae (*Rhizobium*), and Hyphomicrobiales (*Flaviflagellibacter*), except for one isolate of *Fervidobacterium* in the family Fervidobacteriaceae (phylum Thermotogota). Other studies have repeatedly found that Aeromonadaceae, Chitinophagaceae, Pseudomonadacaeae, Sphingomonadaceae, Flavobacteriaceae, and Rhizobiaceae are common members of the core duckweed microbiome (Acosta et al., 2020; O’Brien et al., 2020a,b; Inoue et al., 2022; Baggs et al., 2022).

### Microbial productivity

There was a significant biodiversity effect on microbial productivity. Ten-strain microbial communities were significantly more productive than the average single microbial strain (Table 1, Figure 1), despite starting the experiment at the same cell density. Microbes also grew to significantly higher cell densities in the presence than in the absence of a plant host (Table 1), with slower-growing microbes benefiting most from host presence (Figure 2). Microbial growth without a host significantly predicted microbial growth with a host (Churchill: adjusted *R*^2^ = 0.408, p = 0.020; Wellspring: adjusted *R*^2^ = 0.335, p = 0.036), but strain and community means were always above the 1:1 line (dotted in Figure 2) that would indicate equal growth with and without hosts. Nonetheless, the benefits of microbial diversity and host presence were sub-additive; there was a significantly negative biodiversity x host presence interaction effect (Table 1). The magnitude of this effect indicates that the productivity of 10-strain microbial communities was increased less by the presence of a host than the productivity of single microbial strains (Table 1, Figure 2). In the model in Table 1, there were also significant random effects of plate (p < 0.001) and bacterial strain (p < 0.001) nested within population, but not well location (edge vs. interior) (p = 0.417), on microbial productivity.

**Table 1.** Model results for the effect of microbial strain diversity (one versus ten strains) on microbial productivity. Intercept is a single microbial strain growing in the absence of a host. Estimates are on a natural log scale.

|  | Estimate | SE | df | t-value | p-value |
| --- | --- | --- | --- | --- | --- |
| Intercept | 5.464 | 0.211 | 18.340 | 25.870 | < 0.001 |
| 10-strain community | 2.612 | 0.626 | 22.373 | 4.170 | < 0.001 |
| Host | 2.164 | 0.101 | 404.398 | 21.398 | < 0.001 |
| 10-strain community x host | -1.626 | 0.367 | 397.776 | -4.431 | < 0.001 |

**Figure 1.**
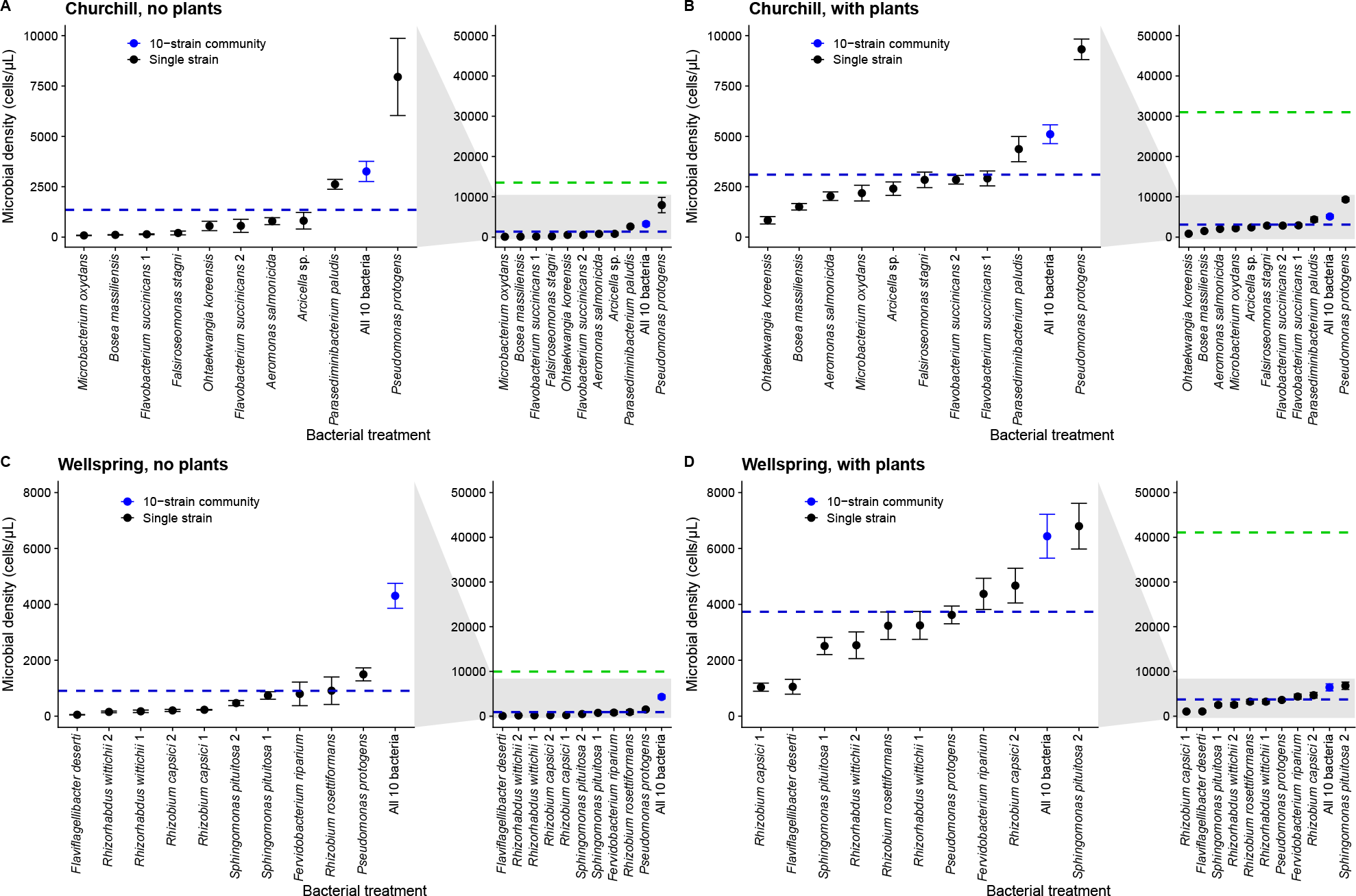
Microbial cell density (±1SE) of bacterial strains and 10-strain synthetic communities in the absence (left) and presence (right) of *Lemna minor* from Churchill (A,B) and Wellspring (C,D). The blue dashed line is the mean of the single-strain effects. The green dashed line is the additive expectation for the 10-strain community.

**Figure 2.**
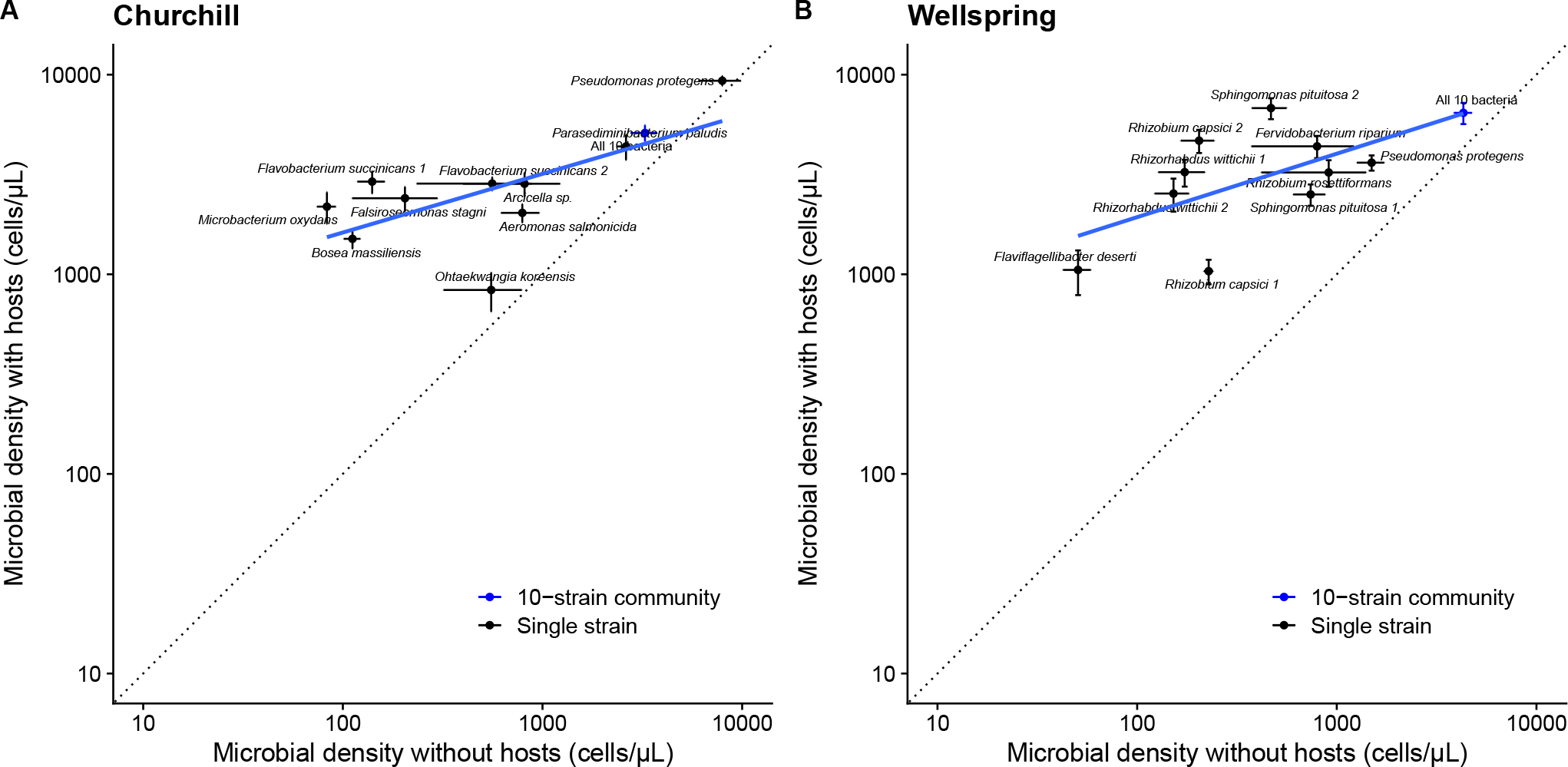
Microbial cell density (±1SE) of bacterial strains (black dots) and 10-strain synthetic communities (blue dots) in the absence (x-axis) and presence (y-axis) of host plants from Churchill (A) and Wellspring (B). Points along 1:1 line (dotted) would indicate microbes or microbial communities that grew equally well with and without a host. Blue solid lines are simple linear regressions.

Microbe-microbe interactions affected microbial productivity in both the presence and absence of hosts. We used estimated marginal means from linear mixed models to calculate the additive expectation for the productivity of 10-strain communities from the single-strain effects (Tables S1, S2). Ten-strain synthetic communities grew faster than almost all of the single strains considered individually, but the productivity of the 10-strain communities was significantly less than the sum of the cell densities of the 10 strains grown separately (compare the blue points and green dashed lines in Figure 1, and see Tables S1, S2). In no case did the 10-strain community achieve a cell density higher than the sum of the single-strain abundances at the end of the experiment (Figure 1), meaning interactions among microbes in 10-strain communities reduced total microbial growth, consistent with widespread competition. Furthermore, the productivity of the 10-strain communities was farther from the additive expectation in the presence (Figure 1B,D) than in the absence (Figure 1A,C) of a host, contrary to our *a priori* expectation that there would be more positive microbe-microbe effects with than without hosts. Instead, the greater sub-additivity when hosts are present suggests greater competition among microbes in the host than in the free-living environment.

### Host growth

There was a significantly positive microbial diversity effect on host growth. Ten-strain microbial communities significantly increased host growth more than the no-microbe control treatment (Table 2, Tukey’s posthoc test: p = 0.042) and more than the average single microbial strain (Tukey’s posthoc test: p = 0.036, Figure 3). The random effects of edge (p < 0.001), plate number (p < 0.001), and population (p < 0.001), but not bacterial strain nested in population (p = 0.794), were also significant in the mixed model shown in Table 2. In contrast, although single bacterial strains had effects on duckweed growth that ranged from weakly negative to positive in Churchill and from neutral to positive in Wellspring (Figure 3), the average single strain did not change plant growth rate relative to the uninoculated control (Table 2, Tukey’s posthoc test: p = 0.682). Of the 20 single strains we tested, only *Pseudomonas protegens* significantly increased the growth of Churchill duckweeds compared to the uninoculated control (linear mixed model: *P. protegens*, p = 0.007, all other strains, p > 0.05).

**Table 2.** Model results for the effect of microbial strain diversity (0, 1, or 10 strains) on plant growth. Intercept is final frond area (in pixels) in the uninoculated control treatment starting at the mean initial frond area. Initial frond area was centered by subtracting the mean and scaled by dividing by the standard deviation.

|  | Estimate | SE | df | t-value | p-value |
| --- | --- | --- | --- | --- | --- |
| Intercept | 22258.34 | 4725.52 | 1.86 | 4.710 | 0.049 |
| Initial frond area (scaled) | 6756.27 | 433.57 | 456.84 | 15.583 | < 0.001 |
| Single strain | 1362.99 | 1614.19 | 20.01 | 0.844 | 0.408 |
| 10-strain community | 5752.16 | 2203.46 | 20.99 | 2.611 | 0.016 |

**Figure 3.**
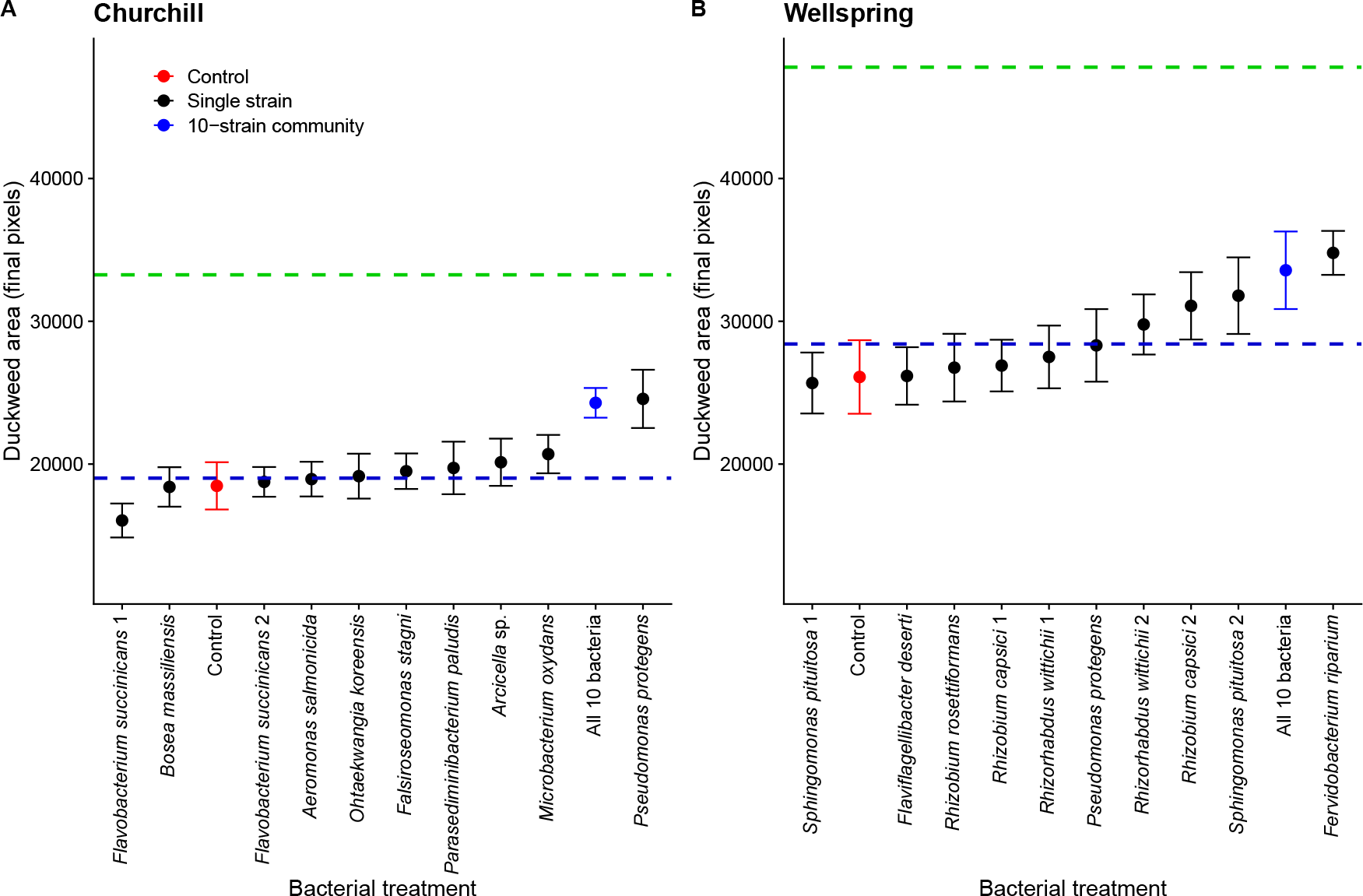
Growth (change in pixel area ±1SE) of *Lemna minor* from A) Churchill Marsh and B) Wellspring Pond in response to bacterial treatment. Red dots are uninoculated control plants, black dots are plants inoculated with a single strain of bacteria, and blue dots are plants inoculated with a 10-strain community. The blue dashed line is the mean of the single-strain effects. The green dashed line is the additive expectation for the 10-strain community. See also Tables S3, S4.

In both populations, the 10-strain community effect was sub-additive and less than the sum of the effects of individual strains on duckweed growth (Figure 3). However, the additive expectation was significantly greater than the effect of the 10-strain community only in Wellspring, and not Churchill (Tables S3, S4); in Churchill, the additive expectation fell just within the upper bound of the 95% confidence interval for the effect of the 10-strain community (Table S3).

### Host-microbe fitness correlations

Duckweed and bacterial fitness were significantly positively correlated in both populations (Churchill: 0.252 ± SE of 0.066, p = 0.004; Wellspring: 0.156 ± 0.058, p = 0.026; Figure 4). No bacterial strains unambiguously benefitted at their hosts’ expense, as would be expected for pathogens.

**Figure 4.**
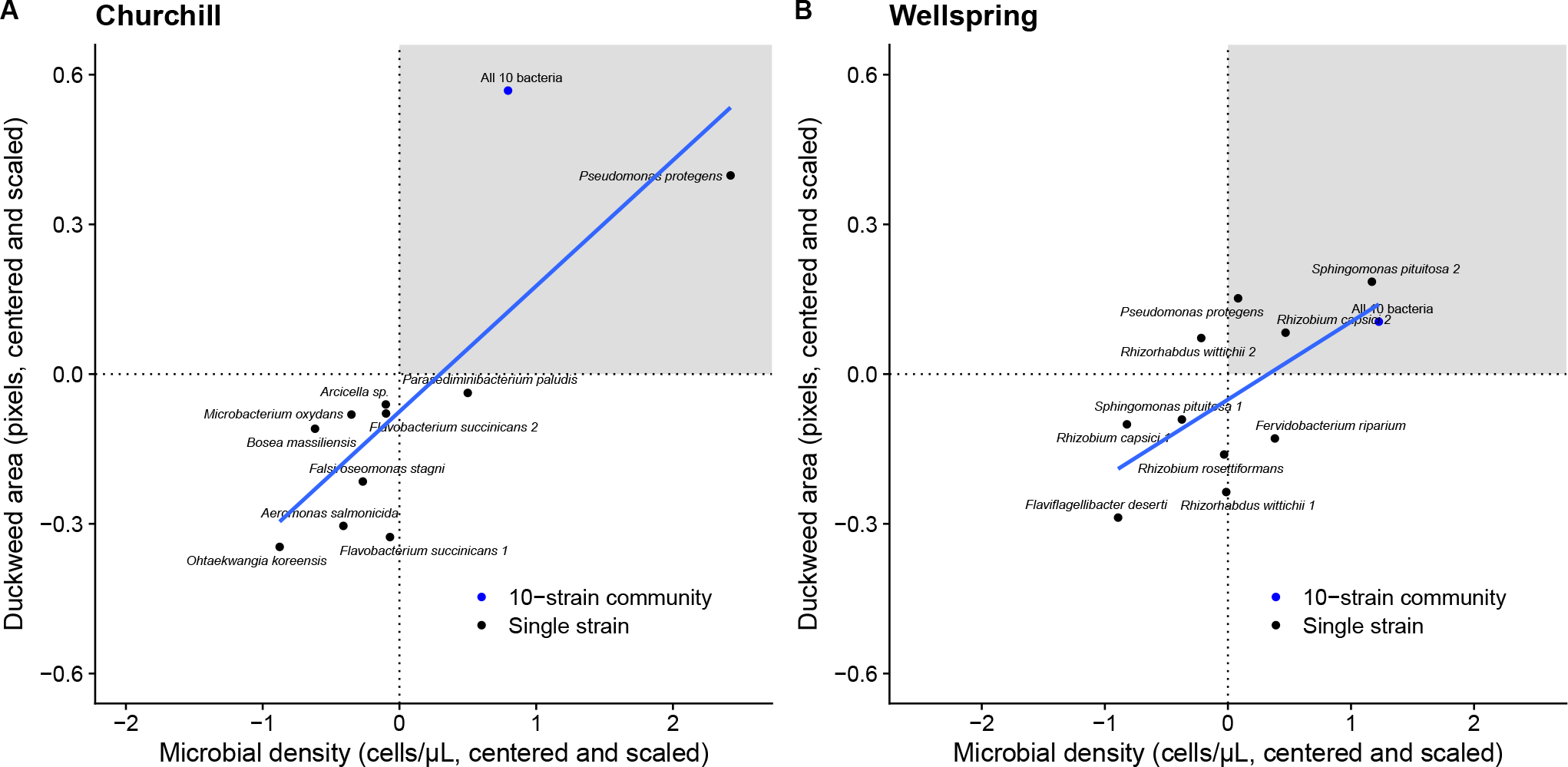
Fitness alignment between bacteria and their *Lemna minor* hosts for A) Churchill Marsh and B) Wellspring Pond. Plotted are estimated marginal means for each single-strain or 10-strain bacterial treatment for duckweed growth and microbial density. The grey shaded regions show strains or communities that achieved above-average cell densities and above-average host growth.

## Discussion

Plant microbiomes are complex communities of interacting microbes, but how interactions among microbes affect the outcome of host-microbiome interactions remains poorly understood. We isolated bacteria that represent many of the dominant taxa in the core duckweed microbiome (Acosta et al. 2020; O’Brien et al. 2020a,b; Inoue et al. 2022; Baggs et al. 2022) from field-collected *L. minor*. We then tested the effects of single strains and 10-strain communities on microbial and host growth. Plant presence sharply increased microbial growth, but most single strains were commensals when tested individually, with only *Pseudomonas protegens* increasing the growth of *L. minor* from Churchill relative to uninoculated controls. However, 10-strain microbial communities led to greater plant growth compared to uninoculated controls, and both microbial and plant productivity were significantly greater with 10-strain communities than in the average single-strain treatment. We found that community effects were generally sub-additive, suggesting that emergent effects of microbiome diversity on microbial and host growth are mostly mediated by competition rather than facilitation among strains. We also tested whether host and microbe fitness interests are aligned and found that the microbes that reached the highest cell densities also provided the greatest benefits to plants.

### Microbial productivity

Microbes grew faster in the presence than the absence of plant hosts, something that is rarely tested in host-microbe experiments. In many systems, the environmental conditions a microbe encounters when free-living are either unknown or hard to recreate in the lab. The microbes that associate with duckweeds, however, live in pond water and thus are readily cultured in minimal media, facilitating direct comparisons of their free-living and host-associated growth rates. Growth in the free-living environment significantly predicted growth with hosts, but not all microbes benefited equally from host presence. The rank order of bacterial productivity sometimes shifted between the host and no-host environment, especially in Wellspring where *Sphingomonas pituitosa* 2 had intermediate productivity in the absence of plants, but was the fastest-growing single strain in the presence of hosts (Figure 1). Furthermore, the strains that grew to the lowest cell densities without a host benefited most from host presence (Figure 2), suggesting their growth is strongly limited by carbon exudates or other host-derived resources.

We found that 10-strain microbial communities were more productive than the average single strain, but effects were always less than the sum of the single-strain cell densities. These results are in line with previous research on BEF and especially diversity-productivity relationships, including results from microbial communities (Tilman et al. 1997a; Bell et al. 2005; Foster and Bell 2012; Delgado-Baquerizo et al. 2016). More diverse communities may simply be more likely to contain ‘keystone’ microbes (such as *P. protegens* in our Churchill population, Figure 1A,B), or diversity-productivity relationships may emerge from niche complementary or facilitation among taxa (Hu et al. 2016; Maron et al. 2018; Piccardi et al. 2019). Our results also match previous findings of mostly sub-additive effects when microbes are co-cultured compared to when they are grown alone (Foster and Bell 2012; Palmer and Foster 2022, and references therein).

Foster and Bell (2012) argued that simple Lotka-Volterra models predict that if two bacteria species compete, the sum of their abundances at equilibrium will be less when grown together than when grown alone. Thus, sub-additive effects such as we observed should be evidence of at least some competition among microbes in mixed communities. Other studies have also employed this framework (e.g., Weiss et al. 2022). According to Foster and Bell (2012), this logic should hold even in substitutive experimental designs such as ours, in which initial cell density was held constant regardless of the number of strains, such that each strain began at 1/10th the cell density in the 10-strain community as in the single-strain treatments. For any given taxon, there were initially ten times more cells in the single-strain treatment than in the 10-strain treatment, but this is unlikely to bias results if microbial growth reached an equilibrial, stationary phase in all treatments (because population size at equilibrium does not depend on initial densities). We did not measure whether microbes had attained a stationary phase, which would have required removing plate seals and risking contamination, but 10 days is more than sufficient for most bacteria to reach their carrying capacities under similar growth conditions as ours. Our results therefore suggest that at least some microbes in our 10-strain communities compete for resources.

Niche overlap is common among microbes, resulting in competition for resources or space. A study of endophytic bacterial communities of the prairie grass *Andropogon gerardii*, for example, found that pairwise combinations of bacteria exhibit a mean niche overlap of roughly 60% (Michalska-Smith et al. 2022). Network and metagenomic analyses of microbial communities (Hester et al. 2019; Xiong et al. 2022) have also underscored the high niche or metabolic overlap among microbes in many microbiomes. Niche overlap is especially likely when microbes are isolated under the same culture conditions, as we did here, and we might have found less competition among strains by isolating microbes on different types of media.

Nonetheless, interactions among microbes in a community are often a mix of competitive and facilitative (Giri et al. 2022), sometimes even within the same pairwise interaction (Venkataram et al. 2023). Members of microbial communities can facilitate one another’s growth despite widespread competition for resources (Piccardi et al. 2019; Giri et al. 2022; Venkataram et al. 2023). Many kinds of mutually beneficial exchanges occur between microbes, such as cross-feeding or complementary ecosystem services such as nitrogen fixation or biofilm formation (Hoek et al. 2016; Goldford et al. 2018; Hassani et al. 2018; Pacheco et al. 2019; Adkins-Jablonsky et al. 2021). For two strains, we can infer that competition is stronger than any mutualistic exchange of metabolites or other services if microbial productivity is less when the two strains are grown together than the sum of their final cell densities when grown apart. However, when comparing the growth of 10 strains in a community to 10 single strains it is challenging to determine how many interactions are competitive versus facilitative. When microbial productivity in a species-rich community does not exceed the sum of the growth of each component microbe in isolation, this merely tells us that competition among microbes is not fully compensated by any microbe-microbe mutualisms that are present; it does not mean facilitation does not occur in the microbiome. Indeed, in the context of limited resources, as the number of species in the community increases, it rapidly becomes impossible for microbial productivity to exceed the sum of individual species effects, as microbial communities reach the maximum density afforded by resources in their environments (e.g., Maron et al. 2018).

We expected to find that hosts increased positive interactions between microbes, because if one microbe increases host growth, it may indirectly benefit another microbe. In our experiments, duckweeds were the only source of fixed carbon available to bacteria, which should result in greater positive interactions among the multiple mutualists of a focal host species (Afkhami et al. 2014, 2021). However, if anything, the productivity of 10-strain communities was closer to the additive expectation from single-strain treatments in the absence than in the presence of plants, suggesting greater facilitation without than with hosts (Figure 1). While this result conflicts with what we expected, it may highlight the ecological contingency of bacterial cooperation on nutrient environment. Several studies (e.g., Piccardi et al. 2019; Zuñiga et al. 2019) have demonstrated that bacterial facilitation is more likely under stressful, nutrient-poor conditions; in our experiments, *L. minor* fronds substantially increased the availability of nutrients in wells through the production of root exudates (Coler and Gunner 1969; Bais et al. 2006), potentially resulting in the emergence of more competitive interactions among strains.

### Host growth

Although single microbial strains benefited from the presence of duckweed hosts (Figures 1, 2), the effect was rarely reciprocal; most single strains had no effect on plant growth, making them commensals when tested in isolation (Figure 3). The only exception was *P. protegens*, which increased the growth of Churchill *L. minor* on its own. *Pseudomonas protegens* is a widespread and often plant growth-promoting bacterium (e.g., Ramette et al. 2011; Narwal et al 2021). However, despite mostly neutral effects of single strains on plant growth, 10-strain synthetic communities significantly increased *L. minor* growth rates, in keeping with other work on the mutualistic effects of microbiomes on duckweeds (O’Brien et al. 2020a,b; Tan et al. 2021; O’Brien et al. 2022, but see Jewell et al. 2023). Thus, beneficial microbiomes can be assembled from strains that are largely commensals in isolation, suggesting that host-microbiome mutualisms are an emergent property of interactions among microbial strains.

The competition among bacteria in 10-strain communities indicated by the microbial productivity results could have either increased or decreased the net benefits microbiomes provide to hosts, depending on whether competitively dominant strains are also highly beneficial to hosts, and on whether the strains they suppress are mainly beneficial or pathogenic for hosts (Afkhami et al. 2014; Hu et al. 2016; Barelli et al. 2020; Löser et al. 2021). Similarly, whether any positive interactions among microbes increase or decrease host benefits depends on whether these interactions promote the growth of microbes that help or harm hosts. Thus, how interactions among microbes affect microbial productivity may not match their effects on host growth.

Nonetheless, we found a similar pattern for 10-strain versus single-strain effects on host growth as for microbial productivity. Inoculation with a 10-strain synthetic community increased host growth more than inoculation with the average single bacterial strain, but did not exceed the benefits of the best-performing single strain. The community inoculation effects we observed may simply reflect the sampling effect, with more diverse communities having higher microbial productivity and host benefits because they are more likely to contain keystone microbes, such as *P. protegens* in Churchill, that exert disproportionate effects on microbial and duckweed growth (Tilman et al. 1997a; Hu et al. 2016; Carlström et al. 2019; Melnyk et al. 2019; Löser et al. 2021; Tan et al. 2021). These results also have implications for microbiome applications in agriculture or medicine; complex communities may not outperform the most beneficial microbial strains.

### Fitness alignment

Outside legume-rhizobium interactions, few studies have measured fitness alignment or conflict between plants and their microbes (but see O’Brien et al. 2020a, 2023). As in O’Brien et al. (2020a), we found that host and microbe fitness were positively correlated in both study populations (Figure 4), although fitness alignment was stronger in Churchill Marsh than in Wellspring Pond. No individual strains significantly increased their own fitness above the population average while reducing host fitness below its population average (Figure 4), as we would expect for strong pathogens or ‘cheaters’ (Jones et al. 2015). The legume-rhizobium literature has also found little evidence of cheaters (Friesen 2012, but see Porter and Simms 2014). Instead, ineffective rhizobia appear to be regularly out-competed and replaced by more beneficial strains (Batstone et al. 2020). That host and microbe fitness interests are closely aligned even in the more facultative associations between duckweeds and microbes is somewhat surprising, given that we did not pre-select only beneficial microbial strains to use in our experiments, and we would have expected to sample some plant pathogens simply by chance.

Compared to the legume-rhizobium symbiosis, plant microbiomes involve much greater bacterial diversity (Dastogeer et al. 2020), and associations between hosts and particular microbes are less reliable across time and space (Jones et al. 2022). Duckweed microbiomes, and plant microbiomes more broadly, are largely environmentally acquired, and while host genotypes can control aspects of microbiome community assembly, such processes often act with low resolution on broad phylogenetic differences among microbes (Fonseca et al. 2018; Acosta et al. 2020; Jacoby et al. 2020; VanWallendael et al. 2022), rather than on the strain-level variation that often mediates microbial effects on plants (Melnyk et al. 2019; Löser et al. 2021; Tan et al. 2021). Nonetheless, our results suggest that duckweed-microbe fitness interests may be be aligned ‘by default’, just as in legume-rhizobium interactions with little coevolutionary history (Friesen 2012). As such, plant-microbiome interactions may not require the evolution of specific host control mechanisms (Kiers et al. 2003; Foster et al. 2017) to maintain mutualism between partners.

## Acknowledgements

We thank Frederickson lab members, especially A. O’Brien and O. Pogoutse, as well as undergraduate students C. Knox, C. Chen, D. Luo, M. Wasim, and others, who helped us troubleshoot and refine our experimental approach. J. Tan helpfully suggested adding a PBS wash to the duckweed sterilization protocol. We acknowledge funding from the Gordon and Betty Moore Foundation (Grant GBMF9356), the Natural Sciences and Engineering Research Council of Canada (NSERC) (Discovery Grant to M.E.F. and CGS-D Alexander Graham Bell Scholarship to J.R.L.), and the University of Toronto.

## Author contributions

JRL and MEF designed the study. JRL and EL collected plants and microbes. JRL cultivated and sterilized plants, and cultured and sequenced microbes. JRL conducted the experiments and collected all data. JRL and MEF analysed data, drafted the manuscript, and finalized the text.

## Data availability

All data and R scripts supporting this manuscript are publicly available at: https://github.com/JasonLaurich/Lemna_single_inoculations.

## Supplementary Information

**Table S1.** Estimated marginal means and 95% confidence intervals (CIs) for the productivity of each single microbial strain and the 10-strain synthetic community from Churchill in the absence (left-hand columns) and presence (right-hand columns) of a host. Estimates are in units of cells/*µ*L.

| Treatment | No host |  |  | Host |  |  |
| --- | --- | --- | --- | --- | --- | --- |
|  | Estimate | 95% CI lower | 95% CI upper | Estimate | 95% CI lower | 95% CI upper |
| *Aeromonas salmonicida* | 687.39 | -674.01 | 2048.78 | 1996.78 | 808.59 | 3184.97 |
| *Arcicella* sp. | 850.24 | -456.00 | 2156.48 | 2757.73 | 1323.88 | 4191.58 |
| *Bosea massiliensis* | 172.96 | -1408.67 | 1754.60 | 1490.98 | 196.35 | 2785.60 |
| *Falsiroseomonas stagni* | 151.04 | -1134.22 | 1436.29 | 2378.78 | 1112.53 | 3645.02 |
| *Flavobacterium succinicans* 1 | 101.90 | -1261.05 | 1464.86 | 2839.56 | 1124.02 | 4555.10 |
| *Flavobacterium succinicans* 2 | 544.47 | -750.33 | 1839.27 | 2791.53 | 1307.78 | 4275.28 |
| *Microbacterium oxydans* | 60.49 | -1264.82 | 1385.80 | 2212.05 | 730.65 | 3693.45 |
| *Ohtaekwangia koreensis* | 471.16 | -964.17 | 1906.49 | 833.18 | -734.54 | 2400.90 |
| *Parasediminibacterium paludis* | 2574.86 | 777.00 | 4372.73 | 4369.78 | 2787.22 | 5952.33 |
| *Pseudomonas protegens* | 7907.95 | 6359.00 | 9456.89 | 9325.02 | 7757.22 | 10892.83 |
| 10-strain community | 3258.23 | 1805.23 | 4711.24 | 5086.94 | 3465.51 | 6708.36 |
| Additive expectation | 13522.45 |  |  | 30995.38 |  |  |

**Table S2.** Estimated marginal means and 95% confidence intervals (CIs) for the productivity of each single microbial strain and the 10-strain synthetic community from Wellspring in the absence (left-hand columns) and presence (right-hand columns) of a host. Estimates are in units of cells/*µ*L.

| Treatment | No host |  |  | Host |  |  |
| --- | --- | --- | --- | --- | --- | --- |
|  | Estimate | 95% CI lower | 95% CI upper | Estimate | 95% CI lower | 95% CI upper |
| *Fervidobacterium riparium* | 950.69 | -569.59 | 2470.96 | 4420.19 | 2898.85 | 5941.53 |
| *Flaviflagellibacter deserti* | 162.92 | -1394.17 | 1720.01 | 1030.22 | -590.03 | 2650.46 |
| *Pseudomonas protegens* | 1530.35 | -101.48 | 3162.18 | 3787.26 | 2286.56 | 5287.95 |
| *Rhizobium capsici* 1 | 172.77 | -1397.62 | 1743.15 | 1153.05 | -336.36 | 2642.46 |
| *Rhizobium capsici* 2 | 139.64 | -1400.50 | 1679.78 | 4785.96 | 3297.26 | 6274.66 |
| *Rhizobium rosettiformans* | 982.80 | -534.05 | 2499.65 | 3454.32 | 1879.12 | 5029.53 |
| *Rhizorhabdus wittichii* 1 | 91.46 | -1385.34 | 1568.26 | 3385.86 | 1838.66 | 4933.06 |
| *Rhizorhabdus wittichii* 2 | 191.66 | -1398.90 | 1782.23 | 2863.33 | 1277.93 | 4448.74 |
| *Sphingomonas pituitosa* 1 | 885.94 | -663.46 | 2435.33 | 2556.60 | 1044.40 | 4068.80 |
| *Sphingomonas pituitosa* 2 | 569.81 | -945.62 | 2085.24 | 6854.24 | 5360.65 | 8347.84 |
| 10-strain community | 4290.43 | 2799.37 | 5781.49 | 6785.64 | 4966.21 | 8605.08 |
| Additive expectation | 9968.46 |  |  | 41076.67 |  |  |

**Table S3.** Estimated marginal means and 95% confidence intervals (CIs) for host growth (final size) with each single microbial strain and the 10-strain synthetic community from Churchill. Estimates are in units of pixels.

| Treatment | Estimate | 95% CI<br>lower | 95% CI<br>upper |
| --- | --- | --- | --- |
| Uninoculated control | 17432.61 | 8614.11 | 26251.10 |
| *Aeromonas salmonicida* | 17249.67 | 8426.63 | 26072.71 |
| *Arcicella* sp. | 19691.11 | 10966.69 | 28415.52 |
| *Bosea massiliensis* | 19047.58 | 10136.60 | 27958.55 |
| *Falsiroseomonas stagni* | 18140.25 | 9229.93 | 27050.58 |
| *Flavobacterium succinicans* 1 | 17119.51 | 8406.84 | 25832.18 |
| *Flavobacterium succinicans* 2 | 19247.03 | 10383.66 | 28110.39 |
| *Microbacterium oxydans* | 19224.18 | 10312.96 | 28135.39 |
| *Ohtaekwangia koreensis* | 16874.00 | 7964.69 | 25783.32 |
| *Parasediminibacterium paludis* | 19698.31 | 10876.70 | 28519.91 |
| *Pseudomonas protogens* | 23856.68 | 15052.57 | 32660.79 |
| 10-strain community | 25307.30 | 16652.41 | 33962.19 |
| Additive expectation | 33254.86 |  |  |

**Table S4.** Estimated marginal means and 95% confidence intervals (CIs) for host growth (final size) with each single microbial strain and the 10-strain synthetic community from Wellspring. Estimates are in units of pixels.

| Treatment | Estimate | 95% CI<br>lower | 95% CI<br>upper |
| --- | --- | --- | --- |
| Uninoculated control | 26257.86 | 18526.25 | 33989.47 |
| *Fervidobacterium riparium* | 27707.19 | 19922.92 | 35491.46 |
| *Flaviflagellibacter deserti* | 25180.41 | 17466.41 | 32894.41 |
| *Pseudomonas protogens* | 31114.63 | 23390.79 | 38838.46 |
| *Rhizobium capsici* 1 | 27565.54 | 19795.61 | 35335.46 |
| *Rhizobium capsici* 2 | 30311.58 | 22593.16 | 38030.01 |
| *Rhizobium rosettiformans* | 26917.00 | 19198.91 | 34635.09 |
| *Rhizorhabdus wittichii* 1 | 25676.23 | 17904.21 | 33448.25 |
| *Rhizorhabdus wittichii* 2 | 30005.09 | 22254.31 | 37755.87 |
| *Sphingomonas pituitosa* 1 | 27742.37 | 20036.76 | 35447.98 |
| *Sphingomonas pituitosa* 2 | 31894.35 | 24206.70 | 39582.01 |
| 10-strain community | 30755.44 | 22986.11 | 38524.78 |
| Additive expectation | 47793.64 |  |  |

